# Decreased integrity of exercise-induced plasma cell free nuclear DNA – negative association with the increased oxidants production by circulating phagocytes

**DOI:** 10.1101/432237

**Authors:** Robert Stawski, Konrad Walczak, Ewelina Perdas, Anna Wlodarczyk, Agata Sarniak, Piotr Kosielski, Pawel Meissner, Tomasz Budlewski, Gianluca Padula, Dariusz Nowak

## Abstract

Strenuous exercise increases circulating cell free DNA (cf DNA) and stimulates blood phagocytes to generate reactive oxygen species (ROS) which may induce DNA strand breaks. We tested whether: (A) elevated cf DNA in response to three repeated bouts of exhaustive exercise has decreased integrity; (B) each bout of exercise increases luminol enhanced whole blood chemiluminescence (LBCL) as a measure of ROS production by polymorphonuclear leukocytes. Eleven men performed three treadmill exercise tests to exhaustion separated by 72 hours of resting. Pre- and post-exercise concentrations and integrity of cf nuclear and mitochondrial DNA (cf n-DNA, cf mt-DNA) and resting and fMLP-stimulated LBCL were determined. Each bout increased concentrations of cf n-DNA by more than 10-times which was accompanied by about 2-times elevated post-exercise rLBCL and fMLP-LBCL. Post-exercise cf n-DNA integrity (integrity index, I_206/78_) decreased after the first (0.59±0.19 vs. 0.48±0.18) and second (0.53±0.14 vs. 0.44±0.17) bout of exercise. There were negative correlations between I_206/78_ and rLBCL (ƍ=−0.37) and I_206/78_ and fMLP-LBCL (ƍ=− 0.40) – analysis of pooled pre- and post-exercise data (n=66). cf-mt DNA integrity (I_218/97_) did not alter in response to exercise. This suggests an involvement of phagocyte ROS in cf n-DNA strand breaks in response to exhaustive exercise.

## Introduction

There are rising evidences suggesting that besides health promoting effects, exhaustive exercises might have potential adverse effects on immune system^1,2^. Therefore, explanation of this issue is essential for training recommendations or customization of individual training load. Interestingly, bouts of strenuous exercise caused rapid increase in cell free DNA (cf DNA) concentration in plasma, moreover it was suggested to be a promising marker of acute exercise induced-metabolic changes in human body^3^. The common process following exhaustive exercises is leukocyte demargination from the vascular, hepatic, pulmonary or spleen reservoirs in parallel with exercise-induced increase in cardiac output and blood flow. This elevates the white blood cell count, however, it seems to be not the main contributor to the exercise-induced increase in cf DNA because the rise in the latter is many times higher than post-exercise leukocytosis^3^. Therefore, mechanisms responsible for this phenomenon could include at least the elevated number of circulating leukocytes accompanied by the increase in their activity leading to release of variety of biomolecules including DNA^3^. This is in line with the recent observation of increased formation of intravascular neutrophil extracellular traps (NETs) that could be responsible for the rise in cf DNA in subjects after exhaustive exercise^4,5^. Respiratory burst of circulating phagocytes (mainly polymorphonuclear – PMNs) in combination with phagocytosis and NETs formation is essential for innate immunity^6,7^. Increased generation of reactive oxygen species (ROS) related to activation of circulating PMNs NADPH oxidase complex was reported in healthy subjects after exhaustive exercise^8,9^. ROS can induce damage to variety of biomolecules including modification of DNA bases and DNA strand breaks. It is possible that simultaneous activation of respiratory burst of circulating PMNs and NETs formation in response to exhaustive exercise can lead to strand breaks of released DNA and thus formation of cf DNA with decreased integrity. cf DNA can be divided in two pools: cell free nuclear DNA (cf n-DNA) and cell free mitochondrial DNA (cf mt-DNA), deriving from nucleus or cytoplasmic mitochondria’s, respectively. Because of lack of nucleosomal structure, cf mt-DNA fragments present different pattern of integrity than cf n-DNA^10^. Moreover, mt-DNA seems to be more sensitive to oxidative stress and other genotoxic damages due to the lack of protective proteins and efficient DNA repair mechanism^11^. Therefore, it seems that integrity of cf mt-DNA could be more affected by ROS increase following exhaustive exercise than that of cf n-DNA. To solve this question, we compared the pre- and post-exercise plasma concentrations and integrity of cf n-DNA and cf mt-DNA in relation to ROS production by circulating PMNs in healthy physically active men subjected to three repeated exhaustive treadmill runs.

## Results

All included men successfully completed the study protocol of three repeated exhaustive treadmill exercises. Run distance was 8.6±5.5 km, 10.7±7.6 km and 10.4±7.2 km for the 1^st^, 2^nd^ and 3^rd^ bout, respectively. Other parameters related to exercise tests, changes of selected markers of muscle damage as well as the metabolic and cardiovascular responses to exercise have been described elsewhere^3^. Each bout induced the increase (p<0.05) in plasma cf n-DNA: from 3.4±1.4 to 38.5±27.5, from 4.1±3.3 to 48.5±26.2, and from 3.1±1.6 to 53.8±39.9 ng/mL after the 1^st^, 2^nd^, and 3^rd^ exercise, respectively. cf mt-DNA increased significantly by about 2- and 2.4-times only after the 2^nd^ (from 229±216 to 450±228 × 10^3^ GE/mL) and 3^rd^ (from 173±120 to 462±314 × 10^3^ GE/mL) exhaustive treadmill run^3^.

### Changes of integrity of circulating cell free nuclear and mitochondrial DNA in response to three repeated bouts of exhaustive treadmill exercise

Pre-exercise integrity of cf n-DNA expressed as I_206/78_ did not alter significantly over the study period. However, the tendency to gradual decrease in I_206/78_ was noted e.g. I_206/78_ = 0.59±0.19 before the 1^st^ bout in comparison to I_206/78_ = 0.50±0.22 before the 3^rd^ one (Table 1). Exhaustive exercise resulted in the decrease in cf n-DNA integrity especially after the 1^st^ (pre-exercise I_206/78_ = 0.59±0.19 vs post-exercise I_206/78_ = 0.48±0.18, p< 0.05) and the 2^nd^ bout (pre-exercise I_206/78_ = 0.53±0.14 vs post-exercise I_206/78_ = 0.44±0.17, p<0.05). In contrast to cf n-DNA integrity, the pre- and post-exercise-integrity of cf mt-DNA (I_218/97_) was relatively stable over the study period and did not alter in response to exercise (Table 1).

**Table 1.** Cell free nuclear (cf n-DNA) and mitochondrial (cf mt-DNA) DNA integrity in eleven average-trained men before and after each of three bouts of exhaustive treadmill exercise.

| DNA Integrity | 1<sup>st</sup> bout (day 7) |  | 2<sup>nd</sup> bout (day 10) |  | 3<sup>st</sup> bout (day 13) |  |
| --- | --- | --- | --- | --- | --- | --- |
|  | Before | After | Before | After | Before | After |
| **cf n-DNA**<br>(I<sub>206/78</sub>) | 0.59±0.19<br>(0.57; 0.36) | 0.48±0.18*<br>(0.46; 0.36) | 0.53±0.14<br>(0.55; 0.13) | 0.44±0.17*<br>(0.46; 0.33) | 0.50±0.22<br>(0.38; 0.36) | 0.45±0.17<br>(0.41; 0.22) |
| **cf mt-DNA**<br>(I<sub>218/97</sub>) | 0.54±0.16<br>(0.59; 0.28) | 0.60±0.26<br>(0.69; 0.39) | 0.63±0.18<br>(0.53; 0.33) | 0.67±0.16<br>(0.63; 0.30) | 0.65±0.18<br>(0.67; 0.16) | 0.66±0.20<br>(0.79; 0.40) |

Eleven average-trained men completed the study composed of four visits at days 1, 7, 10 and 13. After determination of $\text{VO}_2\text{max}$ at day 1, three repeated treadmill exercise tests to exhaustion at speed corresponding to 70% of personal $\text{VO}_2\text{max}$ were executed at days 7, 10 and 13. Results are expressed as mean ± SD (median; IQR). \* vs corresponding value before the bout, $p < 0.05$ .

### Changes of resting and fMLP-stimulated luminol enhanced whole blood chemiluminescence in response to three repeated bouts of exhaustive treadmill exercise

Mean rLBCL increased by about 2.1-, 2.6- and 2.4-times (p<0.05) in response to the 1^st^, 2^nd^ and the 3^rd^ bout of exhaustive exercise (Table 2). The values of pre-exercise rLBCL were relatively stable over the study period and the same was noted for the post-rLBCL. Similarly, behaved mean fMLP-LBCL: 1.7-, 1.8- and 1.5-fold increase (p<0.05) after each exhaustive treadmill run (Table 2). Mutual comparisons of pre-exercise fMLP-LBCL values at consecutive days did not reveal any significant differences. The same was noted for post-exercise fMLP-LBCL. Consequently, the mean ratio of pre-exercise fMLP-LBCL to pre-exercise rLBCL values was similar and ranged from 2.7 at the 1^st^ bout to 3.0 at the 3^rd^ bout. The mean ratio of post-exercise fMLP-LBCL to post-exercise rLBCL was also stable and reached 2.2, 1.8 and 1.9 after the 1^st^, 2^nd^ and 3^rd^ bout of exercise, respectively.

**Table 2.**
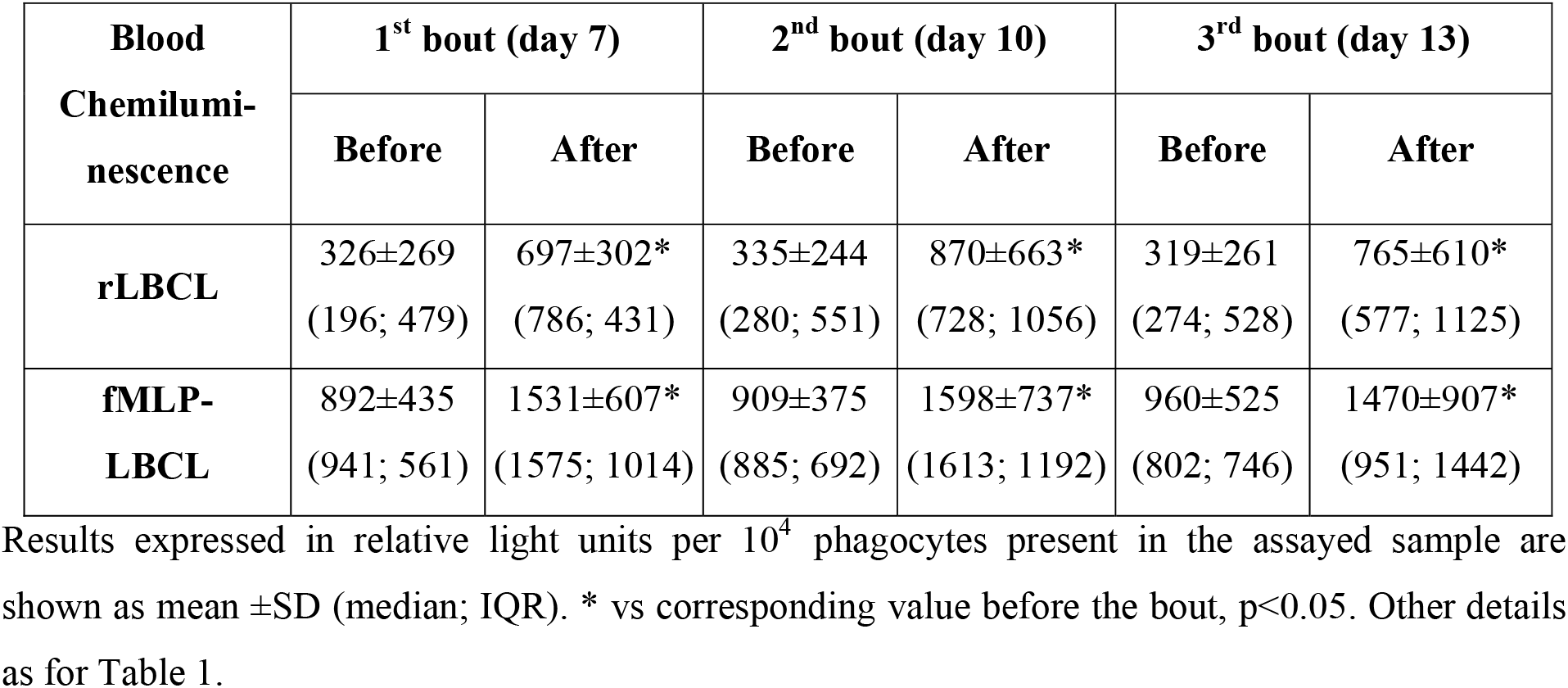
Resting (rLBCL) and fMLP-stimulated luminol enhanced whole blood chemiluminescence (fMLP-LBCL) in eleven average-trained men before and after each of three bouts of exhaustive treadmill exercise.

### Correlations between integrity of cf n-DNA and whole blood chemiluminescence before and after three repeated bouts of exhaustive exercise

Because we studied a relatively low group of men (n=11), the direct analysis of correlations between variables obtained during one bout was inconclusive. Therefore, to overcome (at least partially) this problem, we analyzed correlations between integrity of cf n-DNA (I_206/78_) and LBCL by using the following datasets from three bouts of exhaustive exercise: (A)- pooled individual pre-exercise data (n=33); (B)- pooled individual post-exercise data (n=33); and (C)- pooled pre- and post-exercise data (n=66) (Table 3). There was a negative correlation of moderate strength between pre-exercise I_206/78_ and pre-exercise fMLP-LBCL (ƍ = −0.36, p<0.05), while correlation between post-exercise I_206/78_ and post-exercise rLBCL or fMLP-LBCL reached borderline significance (Table 3). In the case of analysis of dataset involving pre- and post-exercise data together, I_206/78_ negatively correlated with rLBCL (ƍ =−0.37, p<0.05) and fMLP-LBCL (ƍ = −0.40, p<0.05). The same analyses performed between integrity of cf mt-DNA (I_218/97_) and whole blood chemiluminescence (rLBCL, fMLP-LBCL) revealed no significant correlations (ƍ ranged between 0.11 and 0.38).

**Table 3.** Spearman’s (ƍ) correlations between integrity of cell free nuclear DNA expressed as I_206/78_ and resting and fMLP-induced luminol enhanced whole blood chemiluminescence in eleven average trained healthy men before and after three repeated bouts of exhaustive treadmill exercise.

| Correlated variables | rLBCL | fMLP-LBCL | Explanation |
| --- | --- | --- | --- |
| **Integrity of cf n-DNA**<br>($I_{206/78}$) | -0.29 | -0.36* | Pooled individual pre-exercise data from three bouts (n=33) |
| **Integrity of cf n-DNA**<br>($I_{206/78}$) | -0.31# | -0.32† | Pooled individual post-exercise data from three bouts (n=33) |
| **Integrity of cf n-DNA**<br>($I_{206/78}$) | -0.37* | -0.40* | Pooled individual pre- and post-exercise data from three bouts<br>(n=66) |

cf n-DNA – cell free nuclear DNA, rLBCL – resting luminol enhanced whole blood chemiluminescence, fMLP-LBCL – stimulated luminol enhanced whole blood chemiluminescence. Other details as for table 1. \* - $p < 0.05$ , # – borderline significance – $p = 0.08$ , † – borderline significance – $p = 0.06$ .

## Discussion

It is generally believed that regular physical activity has important health promoting effect. However, long term, repeated exhaustive exercises can evoke pathological reactions leading to the development of overtraining syndrome^12–14^. High intensity exercise related to various sports disciplines^15,16^ was reported to cause fast, transient increases in circulating cf DNA reaching concentrations comparable to those observed in trauma or sepsis^17,18^. Some studies suggest that elevated plasma cf DNA levels may by associated with the increased risk of occurrence of overtraining syndrome in athletes^19,20^. Recently, we found that three repeated bouts of exhaustive treadmill exercise induced increase in cf n-DNA and cf mt-DNA in average trained healthy men without development of tolerance^3^. In the present study (which is the continuation of the aforementioned one), we found that the increase in cf n-DNA in response to each of three repeated bouts of exhaustive treadmill exercise is accompanied by the decrease in cf n-DNA integrity. In parallel to this phenomenon, each bout of exercise caused significant increase in rLBCL and fMLP-LBCL and that reflects increased post-exercise ROS production by circulating phagocytes, namely PMNs. Moreover, analysis of pooled data from all bouts revealed a few significant negative correlations between integrity index of cf n-DNA and LBCL. These considerations suggest the involvement of ROS generated from circulating phagocytes in fragmentation of post-exercise cf n-DNA. Although, cf mt-DNA raised in response to the 2^nd^ and 3^rd^ bout of exercise, very surprisingly, no changes of the integrity index of cf mt-DNA were noted over the study period.

### Effect of three repeated bouts of exhaustive treadmill exercise on luminol enhanced whole blood chemiluminescence

Each bout of exercise increased mean rLBCL and fMLP-LBCL in healthy men. Moreover, individual results revealed substantial increase in post-exercise rLBCL and fMLP-LBCL in each volunteer at any occasion. These observations indicate that exhaustive run increased basic generation of ROS by circulating PMNs and primed these cells for increased ROS production after stimulation with fMLP. Our results are consistent with the previous studies showing increased zymosan-induced luminol enhanced chemiluminescence of neutrophils isolated from blood collected after intensive aerobic exercise in healthy men^21–23^.

Moreover, neutrophils of subjects who completed 100 km marathon run revealed increased ROS production as reflected by the afore-mentioned method^24^. On the other hand, other studies did not confirm these observations and even reported suppression of ROS production by neutrophils in the group of well-trained men after marathon run or after acute bout of moderate intensity running and cycling using the same stimulator of phagocyte oxidative burst^25,26^. These differences may result from various study protocols, including exercise load and duration, time of blood collection, usage of whole blood or isolated cells for determination of ROS generation after stimulation with various cell activators. For instance, a continuous 90 min. exercise at the intensity of 50% VO_2_max caused an increase of phorbol 12-myristate 13-acetate (PMA)-induced neutrophils chemiluminescence while zymosan-induced chemiluminescence remained unchanged in healthy men^27^.

PMA is a direct activator of protein kinase C with following activation of NADPH oxidase and neutrophils oxidative burst while stimulation with opsonized zymosan (insoluble cell-wall preparation from the fungi Saccharomyces cerevisiae) involves phagocytosis of these particles, cell degranulation and massive production of ROS^28^. In our study, we used fMLP (an analog of N-formylated bacterial chemotactic peptides) that after binding to specific G protein–coupled receptors on the PMNs plasma membrane activates signal transduction pathways (phospholipase C–dependent generation of diacyl glycerol and inositol 1,4,5-triphosphate, rise in the intracellular Ca^2+^ concentration, protein kinase C activation), leading to the formation of the active NADPH oxidase complex and ROS production^28^. Thus, the peak time – time from agonist addition to appearance of maximal chemiluminescence – is about 7-times shorter for fMLP than for opsonized zymosan^21,29^. This remark may explain the discrepancies between results of our study and some previous investigations^25,26^ about the effect of exercise on ROS production by isolated neutrophils after stimulation with opsonized zymosan.

Moreover, we studied the effect of exercise on LBCL, thus neutrophils and monocytes were not subjected to any process of isolation that could change the cell ability to respond to stimulations with various agonists. Elimination of the risk of cell priming or de-priming during the isolation procedure is the important advantage of this method^30,31^ which allows monitoring of ROS generation by blood phagocytes under conditions that more closely resemble the in vivo situation than the analysis of isolated cells^31,32^. Exhaustive exercise caused essential increase in circulating pro-inflammatory cytokines, namely IL-6, IL-8, granulocyte colony-stimulating factor (G-CSF), macrophage colony-stimulating factor (M-CSF), and granulocyte macrophage colony stimulating factor (GM-CSF)^33,34^. Although these pro-inflammatory cytokines alone induced rather weak oxidative response of PMNs, they strongly enhanced ROS generation in response to secondary stimulation with fMLP^35 – 38^, and that could be an explanation of increased rLBCL and fMLP-LBCL in healthy men after exhaustive exercise.

### Effect of repeated bouts of exhaustive treadmill exercise on the integrity of plasma cell free DNA

Each bout of exhaustive exercise increased several times the concentration of circulating cf n-DNA, and this was accompanied by the decrease in I_206/78_ of post-exercise cf n-DNA. This indicates that pool of cf n-DNA released in response to exhaustive exercise is subjected to factors leading to its enhanced fragmentation. Conversely, no changes in I_218/97_ were noted over the study period, although cf mt-DNA raised significantly after the 2^nd^ and 3^rd^ bout. Thus, one may conclude that exercise-induced release of cf mt-DNA was not accompanied by parallel processes resulting in its additional fragmentation. It is believed that the source of exercise-induced circulating cf n-DNA are NETs^39^. Various factors related to strenuous exercise including heat stress, catecholamines, pro-inflammatory cytokines (e.g IL-6, IL-8) can induce formation of NETs^39^. Intracellular generation of ROS by NADPH oxidase together with their elaboration by myeloperoxidase in intracellular granules are involved in triggering NETs formation^40^. Thus, nuclear DNA mixed with some components of granules before expulsion of NET fibers from PMNs can be exposed to and damaged by ROS, including breaking of DNA strands into shorter pieces. This could be a plausible explanation of increased post-exercise plasma levels of cf n-DNA with decreased integrity and is in line with observed moderate negative correlations between I_206/78_ and rLBCL and fMLP-LBCL. Moreover, previous studies showing increased concentration of 8-oxo-7,8-dihydro-2-deoxyguanosine (an oxidized derivative of deoxyguanosine – marker of cellular oxidative stress and oxidative damage to DNA) in leukocytes and urine of athletes after intense exercise ^41^ support this explanation.

The exercise-induced rise in plasma cf n-DNA is transient^42–44^ probably due to simultaneous increase in the activity of circulating deoxyribonuclease I^44^. Thus, the post-exercise cf n-DNA decreased back to baseline within 0.5 to 2 hours of recovery^44^.

The bouts of exhaustive treadmill exercise were separated by 3 days of rest, and all healthy volunteers did not perform any additional strenuous exercise during the whole 13-day study period. Therefore, one may conclude that the contribution of NETs in maintaining the preexercise cf n-DNA levels was low (and even negligible) and other processes with the unchanging intensities (e.g. necrosis, apoptosis) were the source of baseline pool of circulating cf n-DNA. This may explain the relatively stable concentrations of pre-exercise cf n-DNA and its integrity over the study period. Circulating cf mt-DNA increased in response to exhaustive exercise at the same time as cf n-DNA. However, pre- and post-exercise I_218/97_ of cf mt-DNA did not differ at any occasion. This is quite surprised considering the release of cf mt-DNA from NETs^45^ and formation of NETs from pure mt-DNA in response to skeletal injuries and orthopedic surgery in humans^46^. On the other hand, neutrophils infected with bacteria formed NETs with extracellular fibers containing n-DNA as the main structural component in vitro^7^. It cannot be excluded that neutrophils in response to exhaustive exercise extrude cf n-DNA alone or with very low admixture of cf mt-DNA. Thus, the exercise-induced increase in cf mt-DNA was about 5- to 6-times lower than that of cf n-DNA and no changes of cf mt-DNA integrity were noted.

Acute severe exercise on cycle ergometer until exhaustion increased mitochondrial ROS production in neutrophils of sedentary young males while the same physical exertion had no stimulatory effect on neutrophils mitochondrial ROS after 2 months of regular exercise training program in these subjects^47^.

Aerobic strenuous exercise resulted in an increase in superoxide radical activity in contracting muscles. However, the contribution of muscle mitochondria to this increase was small^48,49^, and even mitochondria generated less ROS during exercise than at rest^48^. We studied average trained healthy men who regularly performed recreational training where some of them participated in sport disciplines in the past. Therefore, it seems that exhaustive treadmill run did not induce increased generation of mitochondrial ROS and subsequent oxidative damage to mt-DNA.

Platelets are able to release microparticles containing functional mitochondria as well as free mitochondria^50,51^. Circulating phospholipase A2 can digest mitochondrial membranes with subsequent release of mt-DNA into extracellular space^50^. Exhaustive exercise (e.g. marathon run) can induce platelets activation and their degranulation^52^. Additionally, short term lifestyle intervention (moderate energy restriction and aerobic training for 6 weeks) increased serum phospholipase A2 activity in overweight or obese subjects^53^.

Consequently, it is possible that platelets could be another source of cf mt-DNA and encapsulating membranes can protect mt-DNA from oxidative attack. On the other hand, even mt-DNA released in response to exercise could be damaged by ROS originated from other sources (e.g. NADPH oxidase), its amount seems to be too little to decrease the integrity of total post-exercise pool of circulating cf mt-DNA. These considerations may also explain no differences between pre- and post-exercise integrity of plasma cf mt-DNA.

There are very scant and unconvincing data on the integrity of exercise-induced cf DNA in humans. Incremental treadmill test until volitional exhaustion induced significant increase in circulating cf n-DNA but without changes of its integrity compared to pre-exercise cf n-DNA in male athletes^54^. On the other hand, 10 km relay race resulted in a significant decrease in the integrity of post-exercise cf n-DNA in a group of recreational runners^55^. Moreover, both these studies did not evaluate circulating concentrations and integrity of pre- and post-exercise cf mt-DNA. Our results showed that repeated exhaustive exercise decreased integrity of circulating cf n-DNA but did not change the integrity of cf mt-DNA.

### Correlations between integrity of cf n-DNA and luminol enhanced whole blood chemiluminescence

Because of the low number of studied volunteers, we pooled pre- and post-exercise data from three consecutive bouts of exhaustive treadmill run and calculated correlations between cf n-DNA integrity and rLBCL and fMLP-LBCL. Such approach has some limitations and can increase the risk of bias, however, for instance, we found significant negative correlations between I_206/78_ and rLBCL, and I_206/78_ and fMLP-LBCL for pooled individual pre- and post-exercise data from three bouts. Moreover, all values of Spearman’s ƍ (n= 6) ranged between −0.40 to −0.29 and that suggests that there is some negative relationship between integrity of cf n-DNA and generation of ROS by circulating phagocytes.

However, according to values of ƍ, these correlations can be interpreted as weak or moderate. This information suggests indirectly that other factors could be responsible for decreased integrity of post-exercise cf n-DNA, or post-exercise LBCL did not reflect precisely the generation of ROS by PMNs during the exhaustive run when NETs were formed and cf n-DNA released. Circulating cf n-DNA increased already after 9 min. from the onset of incremental treadmill run test (at the end of 10 km/h stage corresponding to 63% VO_2_max)^55^. In our study, mean run time was 47 min., 57 min. and 56 min. at the 1^st^, 2^nd^ and 3^rd^ bout, respectively^3^. Thus, pre- and post-exercise LBCL (rLBCL and fMLP-LBCL) reflected the activity of PMNs just before and after the exercise but not during the last 30 min. of run accompanied by the formation of NETs. Moreover, luminol crosses the cellular membrane of phagocytes (PMNs and monocytes); in this way, LBCL can mirror the extra- and intracellular production of ROS^56^.

It is possible that numerous plasma antioxidants can inactivate extracellular ROS (in the close vicinity of PMNs) before their reaction with n-DNA. In consequence, intracellular activity of ROS during formation of NETs seems to be crucial for the decrease of cf n-DNA integrity. These aforementioned reasons may explain the low strength of observed correlations between pooled data of cf n-DNA integrity and LBCL. Concomitant increase in the serum activity of DNase I was recognized as a factor responsible for transient nature of cf n-DNA response to strenuous exercise^57^. Therefore, decreased integrity of post-exercise cf n-DNA could be the result of cleaving NETs DNA by this endonuclease. However, we did not observe any changes of post-exercise cf mt-DNA integrity. For that reason, contribution of DNase I to the decreased integrity of postexercise cf n-DNA seems to be not important under conditions of our study.

### Strengths and weaknesses of the study

Strengths of this study were: (A) the unique protocol consisting of three exhaustive treadmill runs at speed corresponding to 70% of individual VO_2_max separated by 3 days of resting and executed in a standardized laboratory environment; (B) simultaneous measurement of pre- and postexercise concentrations and integrity of circulating cf n-DNA and cf mt-DNA, accompanied by monitoring of resting and fMLP-stimulated LBCL as a marker of ROS production by circulating phagocytes; (C) recruitment of male volunteers (members of our university scientific or administrative staff) highly motivated to comply with instructions related to participation in the study and; (D) exclusion of any subjects (on the basis of medical examination, blood chemistry, spirometry) who could have altered cf DNA levels for reasons other than strenuous exercise. On the other hand, relatively low number of studied subjects (n=11) and exclusion of female volunteers from the trial could be recognized as a weakness of the study. Because this report is the extension of our previous one, all these afore-mentioned limitations were discussed in details elsewhere^3^. In addition, the inhibitory effect of progesterone and estradiol on ROS production by human PMNs in vitro^58^ supported our decision on participation of only male volunteers in the study. Moreover, the low number of studied subjects was counterbalanced by three repeated bouts of exercise. We found that all bouts induced the significant increase in cf n-DNA, rLBCL and fMLP-LBCL, while two bouts were accompanied by the decrease in cf n-DNA integrity and increase in circulating cf mt-DNA without changes of its integrity. These considerations indicate the repeatability of these phenomena despite small studied group of volunteers. However, there are no doubts that further studies involving larger groups of male and female volunteers should be performed especially in the light of analysis of correlation and determination of causality between decreased of post-exercise cf n-DNA integrity and enhanced LBCL.

### Conclusions

We found that repeated bouts of exhaustive exercise separated by three days of resting caused an increase in luminol enhanced whole blood chemiluminescence and in concentrations of circulating cf n-DNA. Post exercise cf n-DNA revealed decreased integrity which negatively correlated with LBCL. Because whole blood chemiluminescence reflects ROS production by circulating phagocytes, one may conclude that oxidants may be involved in the release of cf n-DNA and cf n-DNA strand breaks in response to exhaustive exercise. Although exercise caused moderate increase in plasma levels of cf mt-DNA, its integrity was stable and did not associate with blood chemiluminescence. This suggests a minor role of ROS in exercise-induced changes of cf mt-DNA.

## Methods

### Studied population

The study included eleven apparently healthy, average-trained men (mean age 34.0±5.2 years, mean body mass index 26.2±3.1 kg/m^2^, maximal oxygen consumption VO_2_max 49.6±4.5 ml/kg × min.). The inclusion criteria were: age between 25 and 45 years and a written informed consent before initiating the study procedures. The exclusion criteria were: alcohol and illicit drug abuse, current cigarette smoking, any history of infectious or inflammatory diseases, use of any vitamins or food supplements, or any systemic pharmacological treatment within 3 months prior to the study. All subjects agreed to keep their dietary habits constant during the study period and to comply with the instructions related to participation in the study.

### The study design

The study design was described in details in our previous report^3^. Briefly, the study consisted of 4 visits at the 1^st^, 7^th^, 10^th^ and 13^th^ day of observation. At the first visit (day 1^st^) 11 average-trained men underwent treadmill VO_2_max test and afterwards, at the three consecutive visits (day 7^th^, 10^th^ and 13^th^), participants performed treadmill exercise to volitional exhaustion at speed corresponding to 70% of their personal VO_2_max. Venous blood (15 ml) was collected into vacutainer tubes with EDTA (Becton Dickinson, Franklin Lakes, NJ) within 5 min. before and after each bout of exercise. One milliliter thereof was placed into a separate tube for resting (rLBCL) and n-formyl-methionyl-leucyl-phenylalanine (fMLP) – stimulated LBCL as well as blood cell count (Micros Analyzer OT 45, ABX, Montpellier, France). The rest was used for determination of concentration and integrity of cell free nuclear (cf n-DNA) and mitochondrial DNA (cf mt-DNA). During the whole study period (13 days) volunteers did not perform any exhaustive exercise besides those related to the study protocol. The study was conducted according to the Declaration of Helsinki. The protocol was reviewed and approved by The Medical University of Lodz Ethics Committee (RNN/95/14/KB), and all volunteers provided a written informed consent.

### Blood processing and measurement of cf DNA

EDTA blood samples were centrifuged (1600 × g, 4°C, 10 min.) immediately after collection. The obtained plasma samples were subjected to the second centrifugation (16 000 × g, 4°C, 5 min.) to remove the cell debris and were stored at −80°C for no longer than 4 weeks until isolation of cf DNA with QIAamp DNA Blood Mini Kit (Qiagen GmbH, Hilden, Germany) and measurement of plasma concentrations of cf n-DNA and cf m-DNA with real time quantitative PCR as described previously^3^. Individual results were obtained as a mean from two separate runs and expressed in ng/mL for cf n-DNA and as genome equivalents (GE)/ml plasma (1 GE = 6.6 pg DNA) for cf mt-DNA^59^.

### Determination of plasma cell free DNA integrity

The integrity of circulating cf n-DNA was evaluated by a quantitative real-time PCR (qPCR) targeting the human GAPDH (glyceraldehyde 3-phosphate dehydrogenase) gene (gene ID 2597). The length of the amplicons selected for this assay was 78 and 206 bp, respectively. The ratio of the concentration of the longer amplicon (GAPDH_206_) to the concentration of the shorter one (GAPDH_78_) (ranging from 0 to 1) defined the integrity index 206/78 (I_206/78_), which was used to estimate the fragmentation of cf n-DNA. Higher values of I_206/78_ (e.g. I_206/78_ =1) indicate that all the cf n-DNA molecules are at least 206 bp in length in the GAPDH gene while lower values show that cf n-DNA contains fragments below 206 bp in the same target gene sequence. Similarly two amplicons, one of 218 bp length (encoding part of mitochondrial ATPase 6 gene, ID 4508, and mitochondrial ATPase 8 gene, ID 4509) and the second one of 97 bp length (encoding part of mitochondrial ATPase 8 gene) for calculation of cf mt-DNA integrity index 218/97 (I_218/97_) were chosen. The assay was designed in a way that the forward primer and the probe were the same for each pair of amplicons, whereas two different reverse primers were used. Both, GAPDH_78_ and MTATP_97_ primers and corresponding probes were described and successfully used in previous studies^59^. The primers for longer sequences (GAPDH_206_ primer and MTATP_218_ primer) were planned using *Primer3* software (Table 4). qPCR was carried out in 20 μL of total reaction volume containing 6.5 μL H_2_O, 10 μL TaqMan^®^ Universal PCR Master Mix (Applied Biosystems, Branchburg, New Jersey, USA), 0.25 μL of each of the two matched primers (one forward and one reverse primer for longer amplicon or one forward and one reverse primer for shorter amplicon) (Sigma-Aldrich), 1 μL of a FAM-labeled MT-ATP 8-probe or 1 μL of a MVIC-labeled GAPDH-probe (both probes from Applied Biosystems, Branchburg, New Jersey, USA), and 2 μL of TE buffer containing cf DNA isolated from plasma. The final concentrations of primers and probes were 0.6 μmol/L and 0.4 μmol/L, respectively. Negative control samples received 2 μL of TE buffer without cf DNA from plasma. Reaction was done in duplicate and performed using the 7900 HT Real-time PCR System (Applied Biosystems) under the following conditions: an initiation step at 50°C for 2 min., followed by a first denaturation at 95°C for 10 min., then 40 cycles at 95°C for 15 s. and annealing at 60°C for 1 min. Serial dilutions of human genomic DNA (Roche) (final concentrations from 0.5 ng/mL to 5000 ng/mL) were used to construct the calibration curve (r^2^= 0.9996) for measurement of PCR products.

**Table 4.**
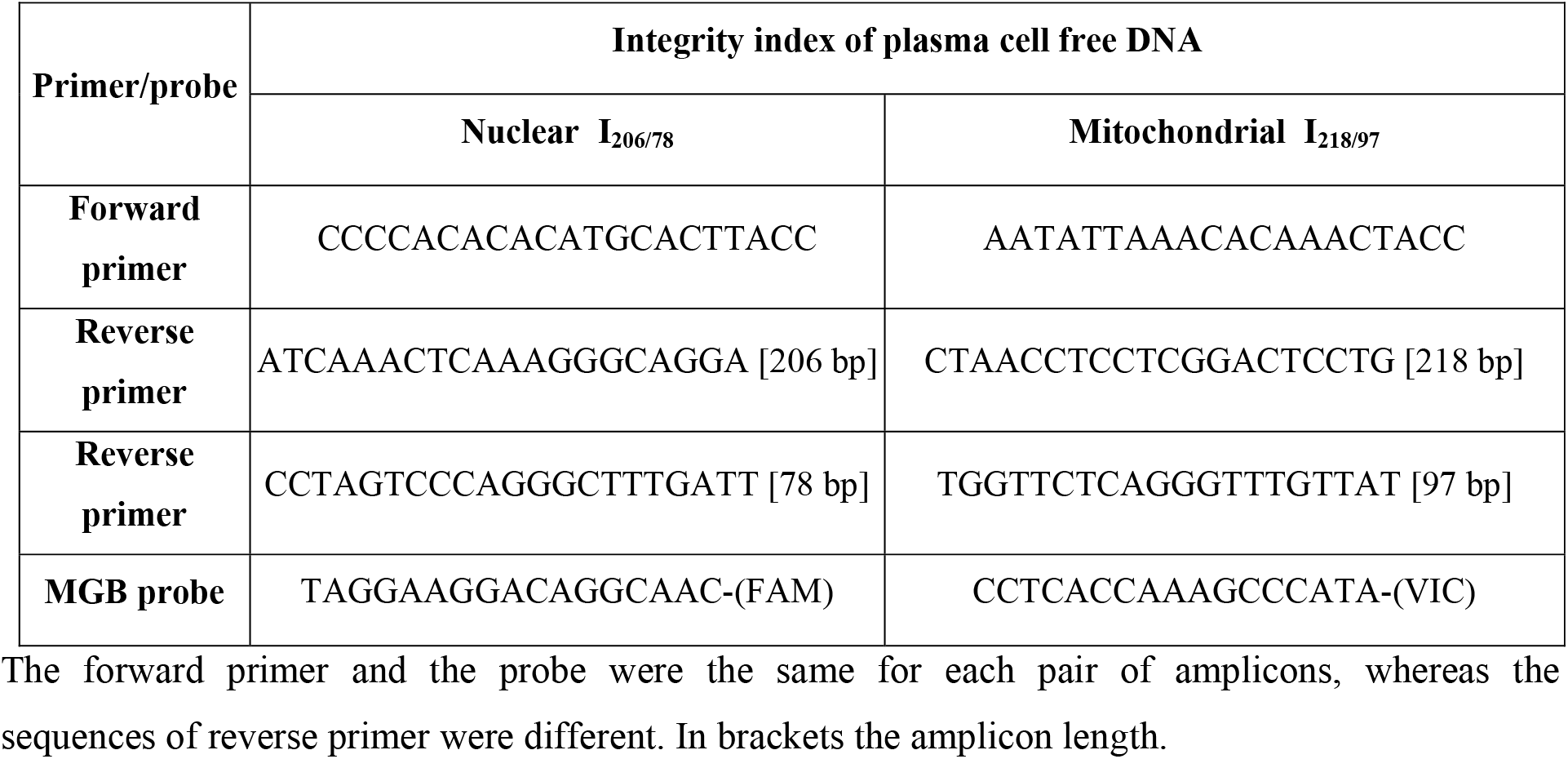
The sequences of primers and probes used for determination of integrity index of circulating cell free nuclear DNA (cf n-DNA) and cell free mitochondrial DNA (cf mt-DNA).

### Resting and fMLP-induced luminol enhanced whole blood chemiluminescence

The resting (rLBCL) and fMLP-induced LBCL (fMLP-LBCL) reflecting ROS production by circulating phagocytes were measured according to the method of Kukovetz et al.^30^ with some modifications^31^. Briefly, 30 μL of venous blood (collected into vacutainer tubes with EDTA) was initially diluted with 1 mL of mixture solution prepared fresh before the assay and containing 127.5 μg/mL of luminol^31^. Then 103 μL of diluted blood was added to 797 μL of mixture solution (to obtain a final blood dilution of 300 times), placed in a multitube luminometer (AutoLumat Plus LB 953, Berthold, Germany) equipped with a Peltier-cooled detector to ensure high sensitivity, and a low and stable background noise signal, and incubated for 15 min. at 37°C in the dark. Then 100 μL of the control solution (solvent for fMLP) or 100 μL of fMLP solution (final agonist concentration in the sample of 2 × 10^−5^ mol/L) for measurement of rLBCL and fMLP-LBCL was added by automatic dispensers and the total light emission was automatically measured for 120 seconds. Individual results were obtained as the mean of triplicate experiments and rLBCL and fMLP-LBCL was expressed in relative light units (RLU) per 10^4^ phagocytes (PMNs and monocytes) present in the assayed sample^31^.

### Statistical analysis

Results are expressed as a mean (SD) and median (interquartile range). Analysis of variance (ANOVA) for repeated observations (parametric test) or Friedman’s ANOVA (non parametric test) was applied for the assessment of changes in variables over time (before and after three consecutive exercise bouts) depending on data distribution which was tested with Shapiro-Wilk’s W test. In case of statistically significant ANOVA, the post hoc analyses were done with Scheffe’s test or post hoc analysis for Friedman’s ANOVA (multiple comparisons at 2 different time points). Correlations between variables were determined using Spearman’s ƍ. A p value<0.05 was considered significant.

## Acknowledgements

This study was supported by a research grant from the Medical University of Lodz No: 503/1-079-01/503-11-001.

## Author Contributions

R.S. D.N. K.W. were involved in conceptualization, study design and methodology, R.S. P.M. collected the data, R.S. E.P. A.S. A.W. conducted experiments, P.K. T.B. K.W. were responsible for participant safety during the execution of treadmill exhaustive exercise,. D.N. G.P. supply resources, R.S. performed data analysis, statistics and visualization, D.N. was responsible for project administration and supervision, R.S. D.N wrote the main text, D.N. R.S. E.P. G.P review and edit the manuscript. All authors approved final submission.

## Additional Information

All data generated or analysed during this study are included in this published article (and its Supplementary Information files).

